# Mechanically Compliant, Precision-Porous Brain Implants Reduce the Foreign Body Reaction and Guide Regeneration

**DOI:** 10.64898/2026.03.24.713981

**Authors:** Ian Dryg, Le Zhen, Rebecca Darrow, Savannah Lawton, Lars Crawford, Robert Robinson, Steve Perlmutter, James D. Bryers, Buddy D. Ratner

## Abstract

Central nervous system diseases and injuries might be treated by implanted devices, tissue regenerative scaffolds, or drug delivery platforms. However, inflammatory CNS responses limit these interventions and may worsen outcomes following damage to the CNS. Via the foreign body reaction, macrophages and glial cells trigger a “glial scar” around implants, reducing device performance, scaffold regenerative ability, or drug delivery potential. Previous studies have shown that stiffness of CNS implants significantly affects glial encapsulation, but few studies have investigated materials that truly match brain tissue stiffness. Porous precision-templated scaffolds with uniform, interconnected, 40 µm spherical pores have shown favorable healing outcomes and a reduced FBR in numerous soft and hard tissue applications. To quantify the effects of both hydrogel compliance (stiffness) and pore size on glial encapsulation, we implanted poly(2-hydroxyethyl methacrylate-co-glycerol methacrylate) (pHEMA/GMA) scaffolds of varying stiffness and pore size for 4 weeks in rat brain. We observed reduced astrocyte encapsulation around PTS compared to solid hydrogel rods, reduced pro-inflammatory macrophage polarization for softer hydrogels versus stiffer hydrogels, and the presence of new blood vessels, neuronal markers and neurogenesis within the pores. Utilizing soft, precision-porous hydrogels could provide a strategy for mitigating glial scarring and improving regeneration in implant-based CNS treatments.

## Introduction

Injuries or diseases of the central nervous system (CNS) might be treated via various therapeutic interventions involving implantable devices, tissue regenerative scaffolds, or drug release systems. Examples of these interventions include neural recording devices for prosthetic limb control [1], scaffolding for spinal cord injury regeneration [2, 3], drug release platforms to treat various diseases [4], or shunts to treat encephalitis.

However, inflammatory CNS responses limit several potential therapeutic interventions and may worsen outcomes following ischemic and traumatic damage to the CNS. Reactive tissue responses that develop around CNS implants are often responsible for time-dependent declines in electrode performance [5–9]. Fibrous encapsulation of scaffolds or drug delivery platforms via the foreign body reaction (FBR) also limit their effectiveness. These tissue responses are largely attributed to activated microglia and astrocytes. Initial tissue damage from implant insertion recruits circulating macrophages, and activates nearby microglia, which have been described as the tissue-resident macrophages of the brain [10]. Activated microglia and macrophages migrate quickly towards the implant where they release cytokines and other signals that activate and recruit astrocytes to the area of the implant. Over the course of a few weeks in rats, the acute stage of the tissue response subsides and gives way to a chronic reactive tissue response around the implant that has been termed “*gliosis*” [11]. This tissue response is analogous to the FBR observed in the CNS. For neural recording devices, the mass of glial cells and deposited extracellular matrix around the device [12, 13] results in an increased electrical impedance [9, 14], and a decrease in recording signal quality over time [6, 7]. In the case of drug delivery platforms, implant encapsulation significantly reduces the ability of released drug to diffuse away from the implant and provide its therapeutic benefit [15]. In the case of hydrocephalus shunts, implant encapsulation has been shown to block fluid flow, making the shunt ineffective [16]. Modulating this neuroinflammatory response around implanted devices could reduce glial encapsulation, allowing for more functional neural recording devices, regenerative scaffolds, and drug delivery platforms. In the current study, we explored a combination of two biomaterial design strategies that have demonstrated reduced implant FBR: *(1)* using precision porous networks, referred to here as porous templated scaffolds (PTS) to reduce FBR, and *(2)* controlling the compliance of the PTS to reduce the mechanical mismatch between implant and brain tissue.

Our research group is developing biomaterial platform technologies to modulate the phenotype of cells that orchestrate inflammation, *i.e.,* microglia, astrocytes, and infiltrating macrophages. Influenced by their microenvironment, macrophages display a spectrum of activation states that enable a wide range of effector activities. Among the phenotypes most studied in the context of FBR, macrophages may become classically activated into a pro-inflammatory phenotype (M1) [17–21], or alternatively activated into one of three phenotypes - M2a, M2b, or M2c [22–27] that are associated with wound healing and regulation of immune responses. An imbalance in the pro-inflammatory vs. pro-healing macrophage populations contributes to the development of chronic inflammation in dysfunctional wound healing environments [28]. Other work suggests that certain mechanical and physical properties of tissue engineered constructs can enhance tissue regeneration by influencing macrophage phenotype [29, 30]. Alongside physical and structural modifications, surface chemistry-based approaches have shown promise for modulating the FBR. By functionalizing biomaterial surfaces with anti-inflammatory peptides, zwitterionic polymers, or targeted neurotrophic factors, researchers can actively attenuate early microglial activation and minimize subsequent astrocytic scarring at the tissue-device interface [31, 32]. Materials that shift the polarization of macrophages towards the pro-healing phenotype may increase the success of remodeling and healing [33–35].

PTS [36] are polymeric constructs comprised of an interconnected porous network with precise, uniform-sized spherical pores and controlled interconnect diameters between pores. Pore diameter, interconnect diameter, and final material geometry can be precisely tuned by modifying fabrication parameters. Originally developed by the National Science Foundation-funded UWEB Engineering Research Center (1996-2007), PTS have been applied in humans [37, 38] (https://ClinicalTrials.gov Identifier: NCT03916731) and are currently available in Europe as a glaucoma shunt developed by iSTAR Medical (Wavre, Belgium). PTS have been fabricated from several polymers, including: poly(2-hydroxyethyl methacrylate) (pHEMA) hydrogel, a biodegradable pHEMA-co-caprolactone (PCL) copolymer, silicone elastomer, fibrin, alginate, and polyurethane; all with pore sizes ranging from 10-160 µm. The regenerative capability of PTS has been studied in a variety of animal models and in human *in vivo* [39–51]. Maximum vascular density and minimum fibrosis are always noted only for 30-40 µm pore diameter PTS and no other pore size. Consistently, 30-40 µm PTS show robust healing, regardless of the material chemistry or the tissue implant site. However, the impact of PTS on neural tissue integration and regeneration has not been evaluated. The work presented here represents the first report on the implantation of PTS into brain tissue. It is expected that similar pore size-dependent effects on healing observed in other tissues will also translate to neural brain tissue regeneration.

Another method shown to be useful for minimizing device encapsulation is to alter the mechanical properties of the implant. Currently available implanted neural interface devices are made of stiff materials such as micro-machined silicon (Utah or Michigan Arrays) or metal microwires. A growing body of evidence shows this stiffness mismatch between silicon/metallic materials (>100 GPa) and brain tissue (∼2-6 kPa) [52] may result in increased reactive tissue responses to neuroprosthetic devices and that reducing the stiffness of neural interface materials may limit adverse tissue responses [53, 54]. However, many studies investigating material stiffness have still featured materials with stiffnesses that are orders of magnitude greater than the stiffness of the brain. In line with the field’s recent focus on soft implantable devices [55], we designed our implants to match the stiffness of brain tissue to ameliorate mechanical mismatch-induced inflammation, thereby reducing glial scar severity. In the study presented here, a panel of PTS materials that varied both PTS pore size and hydrogel stiffness were implanted into rat brain tissue to quantify the subsequent FBR.

The base chemistry of PTS materials was a co-polymeric hydrogel composed of a 1:1 ratio of monomers 2-hydroxyethyl methacrylate (HEMA) and glycerol monomethacrylate (GMA) that allowed stiffness tunability by the varying water content of the pre-polymer mixture [56]. From prior literature, we expect there will be a correlation between polymer stiffness and extent of glial scarring. In addition, we hypothesize that, compared to nonporous implants and PTS with 100 µm pores, 40 µm PTS will achieve the best biointegration. This work represents the first investigation of PTS for regenerative interventions in the CNS. Here we demonstrate that the strategic combination of precision porous architecture and mechanical property (1) substantially reduces glial scarring, (2) modulates macrophage-induced neuroinflammation, and (3) creates an environment conducive to angiogenesis and potential neurogenesis.

## Methods

### Polymer Implant Fabrication

The fabrication procedure for PTS is described in detail previously [40,56]. Briefly, un-crosslinked poly(methyl methacrylate) (PMMA) microbeads with uniform sizes (40 µm vs. 100 µm) are filled into a mold, typically between two glass microscope slides separated by a 1mm thick Teflon gasket. The beads within the mold are sonicated to ensure close packing, then heated to sinter the beads together at their contact points. The heating time and temperature can be varied to precisely control the diameter of the bead fusion connections, which will eventually become the interconnections between adjacent pores in the final scaffold. The empty space between beads in the bead cake is then filled with a monomer solution. To make the pHEMA/GMA co-polymer scaffolds investigated here, the solution components were HEMA and GMA monomers at a 1:1 ratio, the cross-linking agent tetraethyleneglycol dimethacrylate (TEGDMA), the UV initiator Irgacure 651 (2,2-dimethoxy-2-phenylacetophenone), and water. The infiltrated monomer solution was polymerized with a broad-spectrum UV light for 12min (6 mins/side), and then the PMMA beads are solubilized via dichloromethane (DCM reflux), followed by many thorough extraction washes with acetone, ethanol, and water in a graded, sequential manner. The resulting materials are crosslinked hydrogel scaffolds with an interconnected network of spherical pores with precise and uniform diameter.

Using a 50:50 pHEMA/GMA co-polymer, three different pre-polymerization water contents were tested: 50%, 75%, and 85%. Scanning electron micrographs (SEM) of the resulting scaffold structures are shown in **Figure 1a**. The exact recipes are summarized below in **Figure 1b**. Using these formulations, crosslinked polymer scaffolds were fabricated with three different architectures: solid films (not templated), 40 μm pores, and 100 μm pores. By varying water content in the pre-polymer solution, we could control the resulting scaffold stiffness to mimic brain tissue (**Table S1**). Subsequent lyophilization increased scaffold stiffness to enable penetration of scaffolds into the brain surface during implantation. **Figure 1c** shows the compression modulus of scaffolds in a hydrated state and after lyophilization. Compressive Young’s Modulus measurements were taken using an Instron 5543 with 5mm diameter, 1mm thick disks placed between two flat stages and compressed at a rate of 0.1mm/min until 30% compressive strain was reached. Data was collected with Bluehill 2 software, and Compressive Young’s Modulus values were taken from stress-strain curves at 12-15% strain [57]. Solid films did not use bead templating, so there are no templated pores. The higher water content solid films did show some amount of micro-porosity in the lyophilized SEMs of material cross-sections, although those pores (<30 μm for 75% water, < 50 μm for 85% water) were not interconnected and in histology images there were no cells inside any of those pores. It is unknown whether this microporosity exists when polymers are hydrated, or if they only arise after lyophilization.

**Figure 1.**
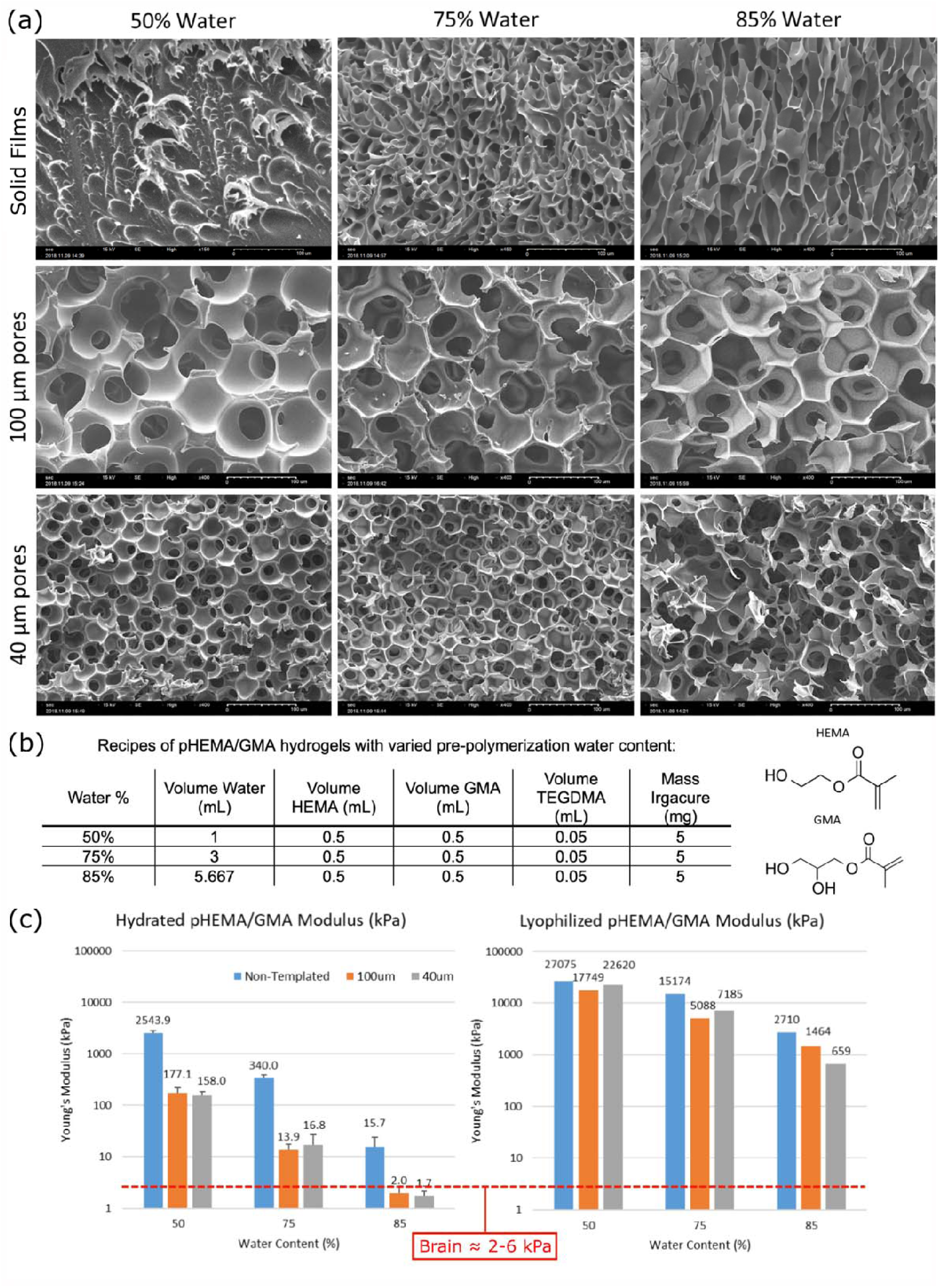
- Characterization of implant materials. **(a)** Scanning electron micrographs of pHEMA/GMA scaffolds with varying pre-polymer solution water content and porosity; scale bars are all 100 μm. **(b)** Recipes of pHEMA/GMA hydrogels with varied pre-polymerization water content. **(c)** Hydrated (left, N=3) and lyophilized (right, N=1) bulk mechanical properties as a function of water content in the pre-polymerization solution and porosity. In lyophilized form, most implants can penetrate brain tissue and then hydrate to soften. Data are presented as mean ± SD when applicable.

After fabrication, a 23 G needle (337 μm ID/641 μm OD) was used to punch out implants with a rod-shaped geometry. To ensure that rods from all polymer groups could penetrate rat brain tissue, they were tested using a 0.83% agar gel insertion test, which has similar mechanical properties to rat pia [58]. Because the softest (85% water) implants were unable to pass the agar brain insertion tests even in their lyophilized form, implants were double-reinforced by coating PTS with sterile-filtered 15% gelatin solution. Gelatin was used because it is not toxic to cells and will quickly dissolve upon implantation into tissue. First, entire slide-sized scaffolds prior to cutting were soaked in 15% gelatin solution at 80°C. The gel was allowed to set at 4°C, and then rods were punched from the scaffolds using the needle punching method. These rods were frozen and lyophilized. A secondary reinforcement was applied by dip-coating the lyophilized polymer rods 1cm deep into sterile-filtered 80°C 15% gelatin, and drying, freezing, and lyophilizing again. Porous polymer film segments left over after punching were also treated in the same way and used for endotoxin testing (Pierce Chromogenic Endotoxin Quant Kit, Thermo Scientific, A39552S). This gel reinforcement procedure was used for all water content groups, including 50% water, 75% water, and 85% water. A no-gelatin control group was included for the 50% water content group to test if there was an effect of the gelatin on the tissue response (this group was chosen because the 50% water group was consistently stiff enough to penetrate the agar gel model). The no-gelatin control group is denoted as g-, and the gelatin-coated groups are denoted as g+. A complete list of the twelve implant groups is: solid 50% water g-, 40 μm porous 50% water g-, 100 μm porous 50% water g-, solid 50% water g+, 40 μm porous 50% water g+, 100 μm porous 50% water g+, solid 75% water g+, 40 μm porous 75% water g+, 100 μm porous 75% water g+, solid 85% water g+, 40 μm porous 85% water g+, 100 μm porous 85% water g+. Only the nine gelatin-reinforced implant groups are included in quantitative analysis in this study.

### Hydrogel Implantation Procedure

All implantations were performed adhering to the approved University of Washington IACUC protocols. Eight young adult female Sprague-Dawley rats (average weight 270g, approximately 15 weeks old) were each implanted with all twelve polymer implants – six per hemisphere in randomly assigned positions. Each animal received general anesthesia (isoflurane 5% induction, 2.5% maintenance). Once induced to a surgical plane, each animal received subcutaneous injections of 10 mg/kg Baytril, 1 mg/kg meloxicam, 0.5mg/kg dexamethasone, and fluids (Lactate Ringer’s solution). A local anesthetic was injected intradermally at the incision site (lidocaine, 1%; bupivacaine, 0.25%). Each animal’s temperature, heart rate, and breathing were continuously monitored during surgery using the SomnoSuite anesthesia monitoring system (Kent Scientific). Fluids were given as needed throughout the surgery. A midline incision was made, and tissue was cleared from the skull’s surface. Bone screws were placed around the perimeter of the opening, and two craniotomies were performed – one over each hemisphere, from ∼2 mm lateral to midline to ∼7 mm lateral to midline, extending ∼8 mm long between Lambda and Bregma. After craniotomies were performed, the dura was opened using fine forceps, and polymer rods were implanted 1cm deep (with tissue at depths of 1-1.5mm imaged and analyzed) with order and position randomized. Craniotomies were covered with gel foam and a PMMA headcap was built covering the exposed skull, stabilized by the bone screws. Post-surgery, each animal received medicated water (Baytril, 0.2mg/mL) for up to 2 weeks.

### Tissue Processing and Histological Analysis

After 4 weeks, animals were injected with Beuthanasia-D, and cardiac perfusions were performed with PBS (to wash out blood), and 4% paraformaldehyde (PFA, to fix the tissues). Carefully, rat brains were removed while ensuring that implanted polymers remained *in situ*. The brains were post-fixed in 4% PFA overnight and then washed in Hank’s buffered HEPES saline (HBHS). Brains were sectioned and sliced using a vibratome into 50-µm thick horizontal slices to display implant cross-sections. Slices were stored in 48-well plates in HBHS with sodium azide at 4° C for storage until staining. Slices from implantation depths of 1-1.5 mm were analyzed, stained with the antibodies shown in **Table S2** for immunofluorescence imaging. GFAP (Glial Fibrillary Acid Protein, astrocytes) and iba1 (ionized calcium-binding adaptor molecule 1, microglia) were used to assess effects on glial scarring; NeuN (neuronal cell bodies), MAP2 (Microtubule-associated Protein 2, dendrites), and Neurofilament (axons) were used to visualize neurons within and around implants; Doublecortin was used to visualize potential neurogenesis; RECA1 (Rat Endothelial Cell Antigen 1) was used to assess vascularization/tissue remodeling; and CD68 (a pan macrophage marker), iNOS (pro-inflammatory macrophages), and Arg1 (pro-healing macrophages) were used to assess macrophage phenotype within and around implants.

Images were captured using a Zeiss LSM 510 confocal laser scanning microscope using 3x3 tiled scans with a Zeiss Plan-Apochromat 20X/0.75 objective lens. All samples used the same antibodies and the same microscope settings (**Table S2**). Image analysis was performed using custom-made scripts in FIJI (ImageJ). For glial cells, fluorescence line profiles were measured. The center point of each implant was selected, then fluorescence profile lines were captured by extending lines radially outwards every 10 degrees (for a total of 36 lines per implant), using the Radial Profile Angle plugin. For each line, the distance from the implant center to the implant edge was noted, so fluorescence profiles could be shifted to start at the implant edge. Once aligned to the implant edge, all fluorescence profile lines from a given implant were averaged. The area under the averaged fluorescence profile from 0-25 μm away from the implant surface was calculated and used as the measure of fluorescence directly surrounding the implant. Astrocyte and microglia fluorescence measurements were compared to control tissue greater than 1 mm from any implants. Even though no implant were present in these control images, to apply the same measurement method we assumed a 337μm diameter faux-implant were present in the control images (because this was the size of the implant fabrication punch). A custom-made FIJI script was developed to assist in cell counting for neuronal density measures and for CD68+, iNOS+ and Arg1+ cell counts. This script allows the user to define the outline of each implant, and then creates 50 μm (for NeuN) or 100 μm (CD68, iNOS and Arg1) thick bands extending from the implant edge. The number of cells within each band was counted using the Cell Counter function in ImageJ. For neurons, cell density was calculated by dividing the number of neuron cell bodies by the area within each band. Neuron cell density measurements were compared to control tissue greater than 1 mm from any implants in the same tissue slice. For CD68+ cell polarization, the iNOS+ (M1) or Arg1+(M2) percent of CD68+ was calculated by dividing the number of iNOS+ CD68+ cells by the number of CD68+ cells, or Arg1+ CD68+ cells by the number of CD68+ cells, respectively. CD68+ cells were characterized in a region 0-100 μm outside of implants.

### Statistical Analysis

Data are presented as mean ± standard error of the mean (SEM). The sample size was n=8 for biological replicates, with one of each of the 9 implant types per animal. For each measure, two-way ANOVA with alpha=0.05 was performed using the statsmodels package in Python (statsmodels.formula.api ols and statsmodels.api.stats.anova_lm) to determine the effects of implant stiffness, implant porosity, and interaction effects. If significant, Tukey HSD pairwise comparisons (statsmodels.stats.multicomp pairwise_tukeyhsd) were performed to determine which groups were significantly different. The Stat annotations Python package was used with Matplotlib and Seaborn to visualize statistical annotations on plots [59]. A star (*) indicates 0.01 < p < 0.05 for Tukey comparisons shown in figures.

## Results

Implant biointegration and inflammatory response were improved by introducing porosity and reducing implant stiffness. Neither porosity nor implant stiffness significantly affected macrophage counts, pro-healing macrophage polarization, microglial encapsulation, or neuron density surrounding implants. However, implant porosity and stiffness affected pro-inflammatory macrophage polarization and astrocyte encapsulation surrounding implants. Implants with 85% water and 40 μm pores showed significantly reduced GFAP fluorescence surrounding implants (**Figure 2b**) and significantly reduced percentage of M1 (iNOS+CD68+) cells (**Figure 3c**) when compared to 50% water, nonporous implants. Markers of neuronal processes and vasculature were observed throughout the porous structure. The 50% water gelatin-control group (i.e. without gelatin reinforcement) resulted in no differences from the 50% water gelatin+ group, so they were excluded from plots for clarity. All materials tested had an endotoxin level <0.025 EU/ml, complying with the standard of sterile materials (<0.06 EU/ml) [56].

**Figure 2.**
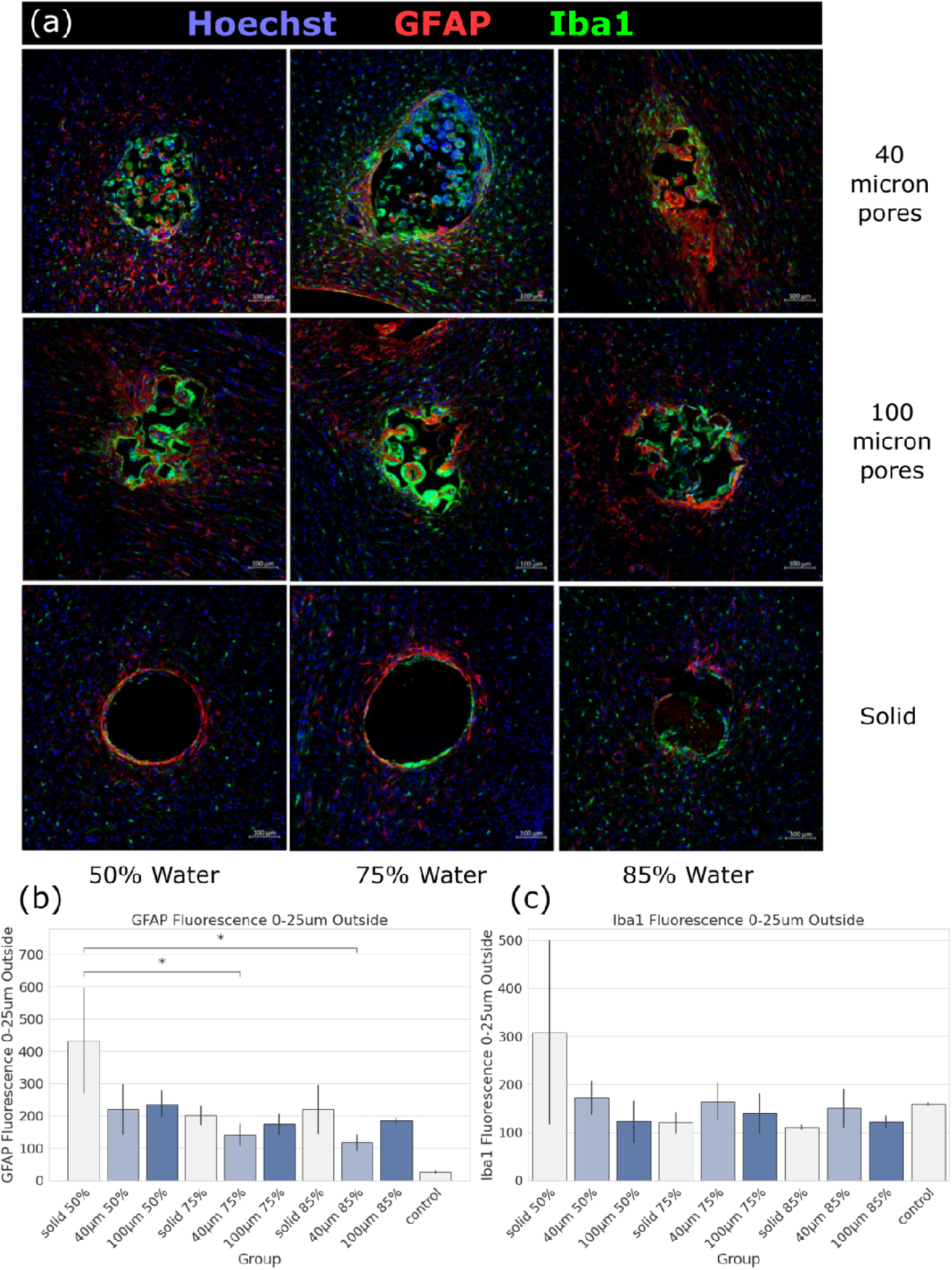
- Confocal microscopy images of glial scars encapsulating implants (cross-sections) for each group (a). Red shows Astrocytes labeled by GFAP, green shows microglia labeled by iba1, and blue shows cell nuclei labeled by Hoechst. Scale bars are 100µm. Effects of polymer water percent and pore size on glial encapsulation of implants is shown for astrocytes via GFAP fluorescence (b), and microglia via iba1 fluorescence (c). Control fluorescence measurements were taken from the same tissue slice, greater than 1 mm from any implants. Significance by post-ANOVA Tukey testing is indicated by * (0.01<p<0.05). Immunofluorescence images shown were randomly selected to provide representative examples of implant shape, porous structure, and biological variations.

**Figure 3.**
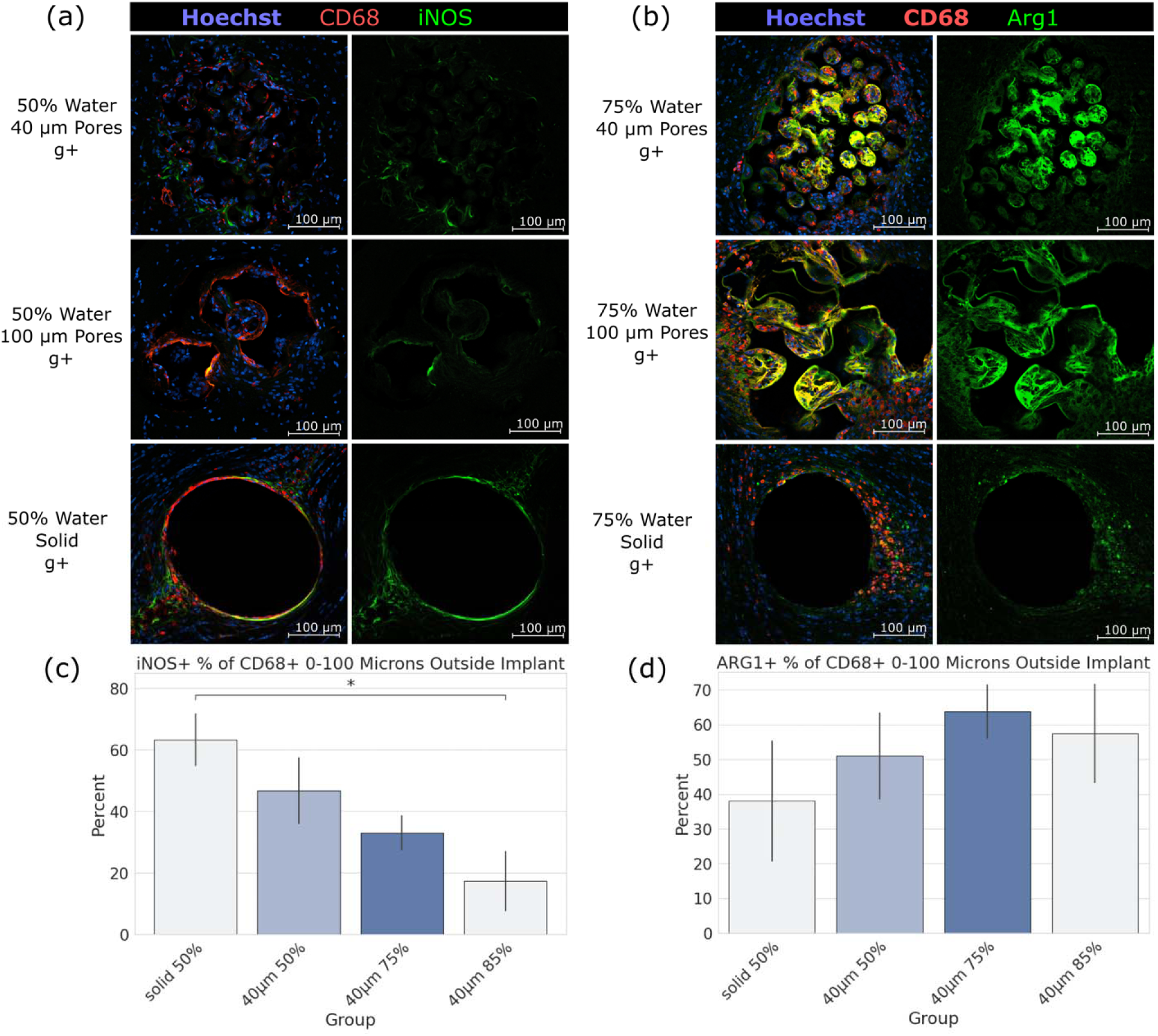
– Implant cross-sections stained to view macrophage polarization: macrophage positivity for iNOS (a,c) and Arg1 (b,d) surrounding implants in green. Red is CD68 (macrophages/microglia), and blue is Hoechst (cell nuclei). Scale bars are 100 μm.

### Soft, porous implants reduce astrocyte encapsulation compared to stiff, solid implants

Considering GFAP fluorescence within 25 μm of implants, material stiffness (p=9.46E-6), porosity (p=4.44E-6), and interaction effects (p=2.86E-5) were all significant by two-way ANOVA (**Figure 2b**). Post-hoc pairwise Tukey HSD comparisons showed that compared to implants with 50% water and no pores, GFAP fluorescence was significantly reduced for implants with 85% water and 40 μm pores (p=0.0152) and for implants with 75%water and 40 μm pores (p=0.0136) (**Figure 2b**). Considering iba1 fluorescence within 25 μm of implants, there were no significant effects, although a trend consistent with GFAP staining was observed.

### Compliant, porous implants improve neuro-inflammation by reducing pro-inflammatory CD68+ cells in comparison to nonporous, stiff implants

Considering percent of CD68+ cells which are iNOS+ within 100 μm of implants, material stiffness (p=6.78E-3) and interaction effects (p=3.63E-2) were significant by two-way ANOVA, but porosity was not. Post-hoc pairwise Tukey HSD comparisons showed implants with 85% water and 40 μm pores significantly reduced iNOS+ percent of CD68+ compared to implants with 50% water and no pores (p=0.0395) (**Figure 3c**). Considering percent of CD68+ which are Arg1+ within 100 μm of implants, there were no significant effects.

### Porous implants promote integration of neural processes throughout the porous network

Considering NeuN+ cell density within 50 μm of implants, there were no significant effects by two-way ANOVA **(Figure 4c).** However, we observed positive labeling of neuronal markers within the pores of PTS, including NeuN+ cells indicated by arrows **(Figures 4b and 5b)**, Neurofilament **(Figure 4a)**, and MAP2 (**Figure 5a**).

**Figure 4.**
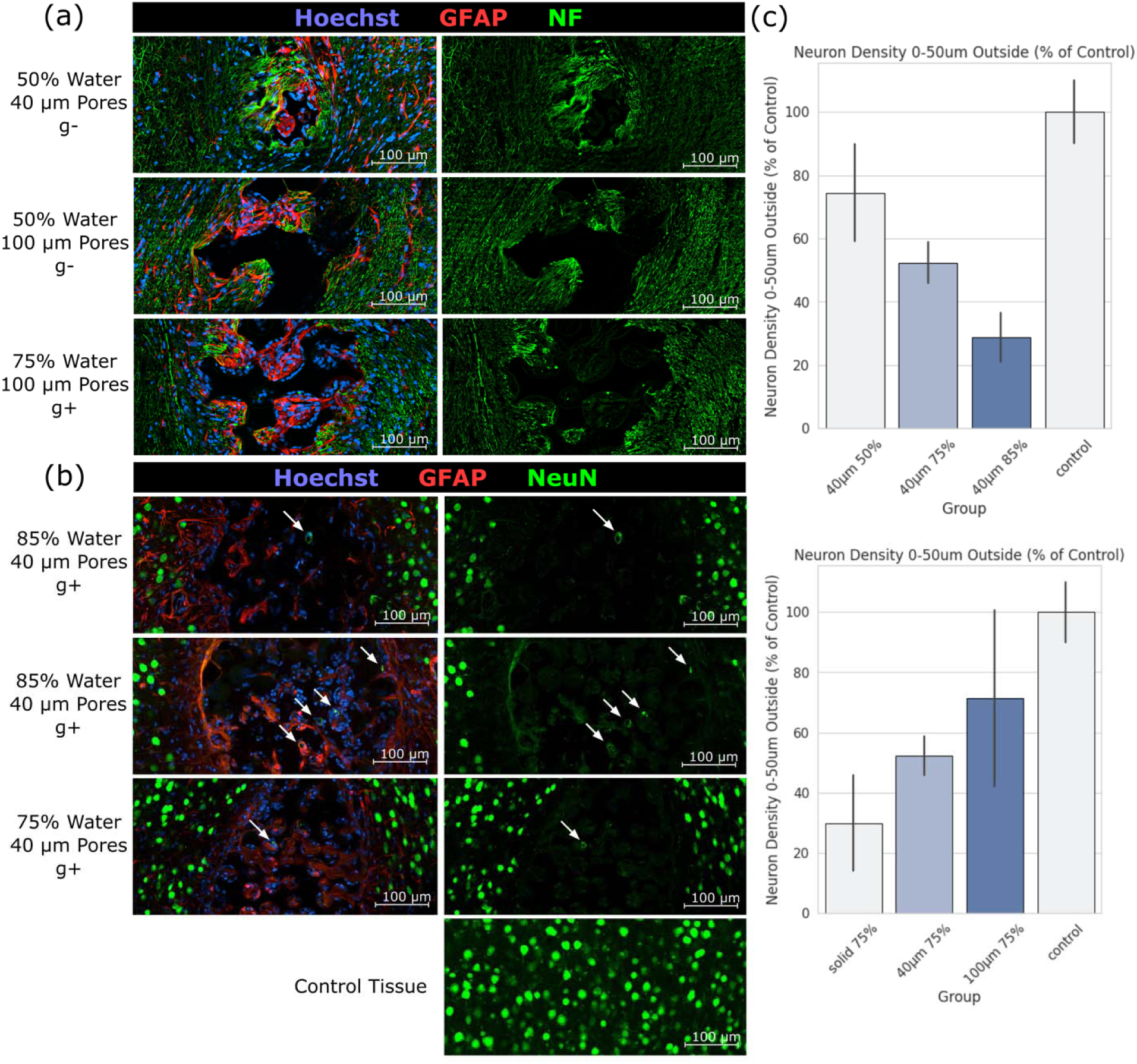
– Implant cross-sections stained to display examples of positive neuronal marker expression around and within pores in green: axons via Neurofilament (a), and neuronal nuclei via NeuN indicated by arrows (b). Blue is Hoechst (cell nuclei) and red is GFAP (astrocytes). Scale bars are 100 μm. NeuN+ cell density surrounding implants is shown (c) for 40 μm and 75% groups. Control NeuN+ cell density measurements were taken from the same tissue slices greater than 1 mm from any implants. Images shown were not selected to match trends in the quantitative data, but to provide representative examples of neuronal processes and cell bodies within the porous structure.

**Figure 5.**
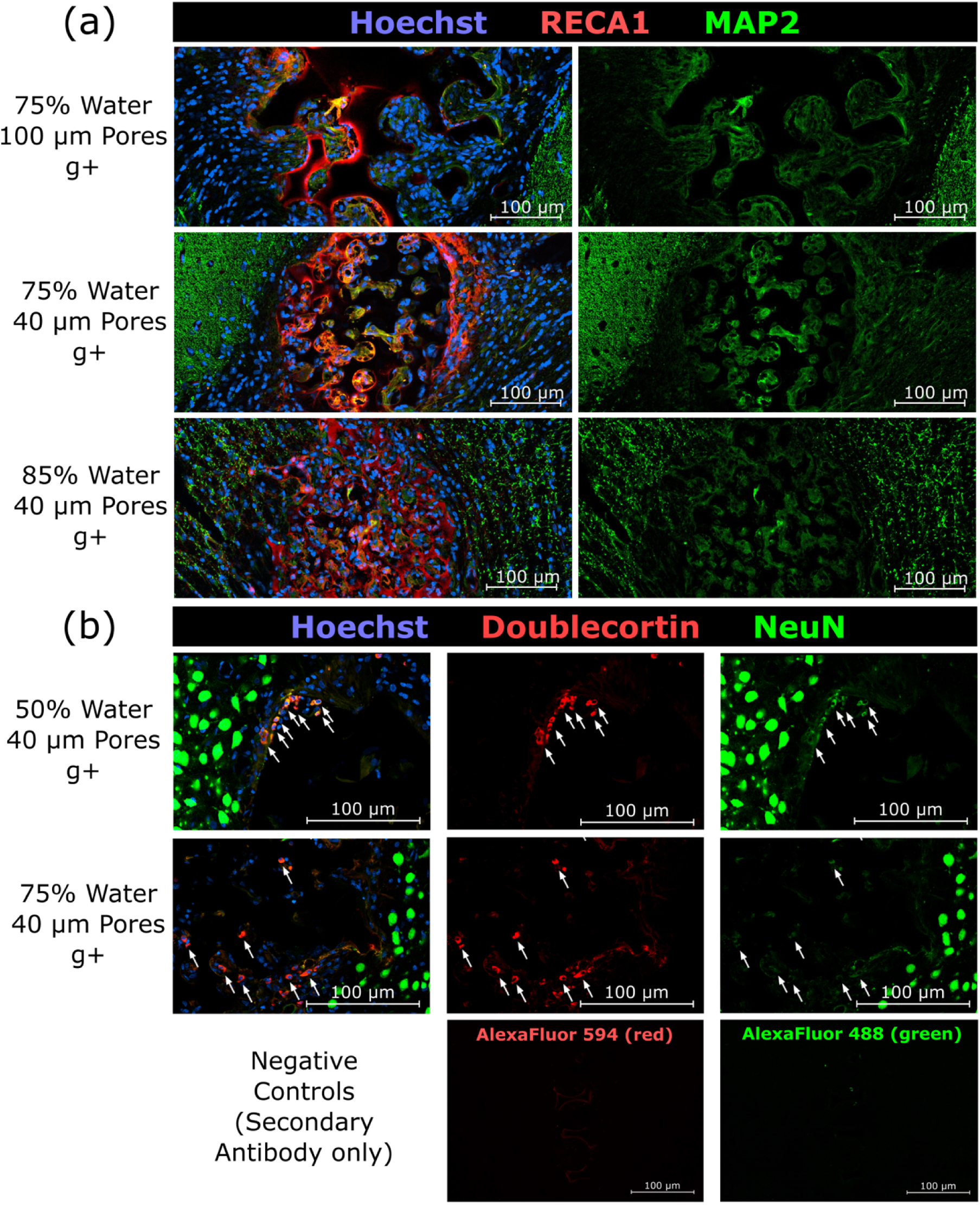
– Implant cross-sections stained to display examples of vasculature marker RECA1 (red) and neuronal dendrite marker MAP2 (green) within pores (a); and new neuron marker Doublecortin (red) colocalized with NeuN+ cells (green) within pores (b). Blue is Hoechst (cell nuclei), scale bars are 100 μm.

### 40 µm PTS promote angiogenesis and neurogenesis

RECA1, a marker for vascular endothelium in rats, labels positive throughout the porous structure indicating blood vessel formation through the pores (**Figure 5a**). In follow-up staining, NeuN+ Doublecortin+ cells were observed within the pores **(Figure 5b)**, suggesting that neurons are proliferating or restructuring [60, 61] to integrate with the porous network.

## Discussion

Our findings demonstrate that the strategic combination of precision porous architecture and mechanical property substantially reduces glial scarring, modulates macrophage-induced neuroinflammation, and creates an environment conducive to angiogenesis and potential neurogenesis. This work represents the first investigation of PTS for regenerative interventions in the CNS.

Excessive, neuro-inhibitory glial scarring characterized by GFAP^+^ astrocytes has long been associated with long-term brain implant failure. In this study, we found that the optimal combination of mechanical property (∼2 kPa) and porous structure (uniform 40 µm spherical pores) significantly reduces glial scarring. Normal neuro-electrodes, for example, have Young’s modulus on the magnitude of hundreds of GPa [53, 54], while the Young’s moduli of CNS tissues range from 2 to 8 kPa [52]. Thus, there is an 8-order-of-magnitude difference in mechanical property between neuro-electrodes and brain tissue. Such a drastic mismatch of mechanical property is likely to exacerbate glial scarring. Indeed, glial scarring has been significantly reduced when the bulk mechanical property of implants was decreased from hundreds of GPa to about 12 MPa, even when the surface mechanical properties are identical [53, 54]. Coating a stiff material with soft hydrogel (11.6kPa by AFM) can also reduce glial scarring [62]. The glial scar mitigation observed in our study is consistent with these previous studies. We further extend previous studies by using a soft hydrogel scaffold that fully matches the stiffness of brain tissue rather than only reducing the stiffness mismatch.

In the study of porous biomaterials, establishment of an optimal pore size remains an ongoing area of research. Previous data provides evidence that 40 µm diameter pores represent an optimal size for healing and biointegration across many tissues. In the current study, pore size was a significant factor in astrocyte encapsulation, and a significant interaction effect for reduction in iNOS+ macrophages, providing a significant role in the biointegration of these implants.

We found many M1 (iNOS+ CD68+) and M2 (Arg1+ CD68+) macrophages residing within the porous structure of the soft hydrogel scaffolds. The percentage of M1 macrophages directly surrounding the implant was significantly reduced for the soft, 40 µm PTS compared to the stiff, nonporous scaffolds. In previous studies, macrophages of diverse phenotypes were observed residing in the porous structure, but those macrophages were a more pro-inflammatory M1 phenotype [40, 63, 64]. The relatively low level of M1 macrophages within the porous structure observed here may be attributed the low stiffness of material. It is important to clarify that M1 response is not necessarily inhibitory to healing, but an integrated part of it.

Under sterile conditions, M1 macrophages produce the highest level of pro-angiogenic and pro-neurogenic signals, vascular endothelial growth factor (VEGF) and fibroblast growth factor 2 (FGF2), compared to M2 phenotypes [65]. This observation could explain why high M1 polarization in pores leads to robust angiogenesis as observed in our studies, where M1 and M2 macrophages coordinate to orchestrate healing. In a transcriptional study of long term implantation of a relatively flexible (∼8 GPa) neuro-electrode [66], it was shown that M1 response dominates in the first two weeks of implantation (corresponding to upregulation of pro-angiogenic genes) and gradually shift towards M2 response after two weeks (a sign of moving towards homeostasis) [67]. The relatively low M1 level within the porous structure observed in this study may indicate an accelerated healing process, or may be due to the fact that tissues were analyzed at 1 month post-implantation.

Perhaps the most striking and unexpected finding in this study was the identification of NeuN+ cells colocalized with doublecortin (DCX+) within the pores of 40 μm PTS. This double-positive labeling pattern is consistent with a transition from immature neuronal precursors to mature neurons [68], suggesting that active neurogenesis may occur [69–71] within the scaffold environment. This finding warrants careful interpretation, as neurogenesis is not typically robust in the neocortex of adult rodents, despite occasional reports of low-level neurogenesis in this region. Neurogenesis is known to mainly occur in two regions of adult rodent brains: the dentate gyrus of the hippocampus [72] and the sub-ventricular zone (SVZ) [73]. Given that our analyzed tissue was at depths of 1-1.5 mm, it is unlikely to have captured these deeper tissue regions. More likely, the scaffolds were in the neocortex, where neurogenesis has occasionally been observed [74, 75].

Injury to brain tissue has been shown to increase neurogenesis around the injury site [76]. The implantation injury itself may have triggered migration of doublecortin+ neuronal precursors from the SVZ to the injury site, as previously seen in mouse cortical lesions [77], where ectopic progenitors migrated along blood vessels and glial cells. Given that both blood vessels and glial cells grow throughout the porous structure during healing, this could explain how precursors could migrate from the SVZ and into the pores. Similarly, doublecortin+ neuroblasts were observed following ischemic injury in mouse cortex, which primarily originated from SOX2+ astrocytes already in the cortex [78], suggesting astrocytes as a source for neuronal trans-differentiation. Another possibility could be that oligodendrocyte precursor cells (OPCs), which maintain dividing cell populations in adult cortex [79] and respond to injury, became reprogrammed into neurons as seen in a recent rat study that turned on transcription factor gene neurogenin2 [80]. Astrocytes in young mice and in SVZ of adult mice can be turned into stem cells *in vitro* by treatment with proangiogenic and pro-neurogenic factors, such as FGF2 [81, 82], and fate mapping shows that astrocytes in young mice can differentiate into neurons during maturation [83]. Endothelial cells can promote trans-differentiation of astrocyte into neural progenitor cells [84], and VEGF also promotes astrocyte-to-neuron trans-differentiation [85]. The macrophages infiltrating our scaffolds might readily produce VEGF and FGF2 [65]. The abundance of endothelial cells and macrophages observed in the scaffolds could be the source of these and other proangiogenic factors. These theories do not fully explain the observations, as doublecortin+ NeuN+ cells are only found within 40 µm pores and not in surrounding tissues, suggesting that the unique immune response created by these scaffolds is required for the observed neurogenesis. These proposed mechanisms are not definitively tested by our current data and represent important avenues for future investigation. The potential for the scaffolds to promote neurogenesis can be further examined in future studies using 5-bromodeoxyuridine labeling (BrdU, marker for DNA replication) [75], Nestin (marker for neural stem cells) [68], and Ki-67 (marker for cell proliferation). The observations from this study set the stage for follow-up studies and new developments for pro-neurogenic biomaterials.

Numerous biomaterials approaches have previously attempted to promote angiogenesis and neurogenesis in adult CNS, via drug delivery or biomaterial structure. Administration of neurotrophic factors, such as Brain-derived Neurotrophic Factor (BDNF) [86] and VEGF [87], have been found to improve neurogenesis in rat stroke models. To stimulate neurogenesis, the stromal cell-derived factor-1a (SDF-1a) was delivered to the stroke site in a targeted fashion by a pH-sensitive polymer [88]. Neurotrophic factors, for example, nanoparticle-clustered VEGF [89], neurotrophin-3 (NT3) [90], and a cocktail of neurotrophic factors and small molecules [71], were packaged with naturally derived hydrogels and injected into damage site in the CNS. In general, the neurotrophic factor delivery approach is expensive and difficult to sustain over long periods of time. Natural polymers, such as collagen [70] and aligned fibrin hydrogel [69] can serve as scaffolds to guide neurogenesis. Aligned synthetic polymer poly-lactic acid (PLA) fibers have shown enhanced neurogenesis in young mice brain [91], but the mechanical property mismatch between the synthetic polymer and the brain was substantial. Overall, our approach builds upon other biomaterials approaches by harnessing the pro-healing effect of PTS, as seen in many non-neuronal tissue sites. Such pro-angiogenic and pro-neurogenic scaffolds have potential for rapid translation to the clinic with broad applications in stroke [88, 89], traumatic brain injury [69, 70, 90, 91], spinal cord injury [68], and even brain aging [92]. Conductive polymer scaffolds may someday provide solutions for next-generation soft neural electrodes [93].

The findings presented here suggest broad potential applications for soft, precision-porous scaffolds in CNS regeneration. These biomaterials could serve as platforms for treatment of acute CNS injuries (traumatic brain injury, stroke, spinal cord injury), chronic neurodegenerative conditions, and potentially as foundations for next-generation brain-computer interfaces or drug delivery platforms. The reduced glial scar formation observed in this study directly addresses a primary limitation of current electrode-based neural interfaces, suggesting potential to extend device functionality. Furthermore, the pro-angiogenic and pro-neurogenic properties of these scaffolds align with therapeutic goal of regeneration across multiple CNS pathologies.

## Acknowledgements

This research was funded by University of Washington Engineered Biomaterials (UWEB) Engineering Research Center, and by University of Washington Institute of Translational Health Sciences Collaboration Innovation Awards.

## References

1. O.A. Mokienko, “Brain–Computer Interfaces with Intracortical Implants for Motor and Communication Functions Compensation: Review of Recent Developments,” Sovremennye Tehnologii v Medicine 16, no. 1 (2024): 78–89. doi: 10.17691/stm2024.16.1.08

2. J. Koffler, W. Zhu, X. Qu, et al., “Biomimetic 3D-printed scaffolds for spinal cord injury repair,” Nature Medicine 25, no. 2 (2019): 263–269. doi: 10.1038/s41591-018-0296-z

3. B. Shrestha, K. Coykendall, Y. Li, et al., “Repair of injured spinal cord using biomaterial scaffolds and stem cells,” Stem Cell Research & Therapy 5, no. 4 (2014): 91. doi: 10.1186/scrt480

4. J. U. Pothupitiya, C. Zheng, and W. M. Saltzman, “Synthetic biodegradable polyesters for implantable controlled-release devices,” Expert Opinion on Drug Delivery 19, no. 10 (2022): 1351–1364. 10.1080/17425247.2022.2131768

5. A. Carnicer-Lombarte, S.T. Chen, G.G. Malliaras, and D.G. Barone, “Foreign Body Reaction to Implanted Biomaterials and Its Impact in Nerve Neuroprosthetics,” Frontiers in Bioengineering and Biotechnology 9 (2021): 622524. doi: 10.3389/fbioe.2021.622524

6. J. C. Barrese, N. Rao, K. Paroo, et al., “Failure mode analysis of silicon-based intracortical microelectrode arrays in non-human primates,” Journal of Neural Engineering 10, no. 6 (2013). 10.1088/1741-2560/10/6/066014

7. M. A. M. Freire, E. Morya, J. Faber, et al., “Comprehensive analysis of tissue preservation and recording quality from chronic multielectrode implants,” PLoS ONE 6, no. 11 (2011): 23–27. 10.1371/journal.pone.0027554

8. A. Prasad, Q. S. Xue, V. Sankar, et al., “Comprehensive characterization of tungsten microwires in chronic neurocortical implants,” *Proceedings of the Annual International Conference of the IEEE Engineering in Medicine and Biology Society*, EMBS, 056015, (2012): 755–758. 10.1109/EMBC.2012.6346041

9. M. P. Ward, P. Rajdev, C. Ellison, and P. P. Irazoqui, “Toward a comparison of microelectrodes for acute and chronic recordings,” Brain Research, 1282 (2009): 183–200. 10.1016/j.brainres.2009.05.052

10. A. Aguzzi, B.A. Barres, and M. L. Bennett, “Microglia: Scapegoat, Saboteur, or Something Else ?” Science 339, no. 6116 (2013): 156–161. 10.1126/science.1227901

11. P. Stice, and J. Muthuswamy, “Assessment of gliosis around moveable implants in the brain,” Journal of Neural Engineering. Aug;6, no.4 (2009): 046004. doi: 10.1088/1741-2560/6/4/046004. Epub 2009 Jun 25. PMID: 19556680; PMCID: PMC2813571.

12. R. Biran, D. C. Martin, and P. A. Tresco, “Neuronal cell loss accompanies the brain tissue response to chronically implanted silicon microelectrode arrays,” Experimental Neurology 195, no. 1 (2005): 115–126. 10.1016/j.expneurol.2005.04.020

13. J. N. Turner, W. Shain, D. H. Szarowski, et al., “Cerebral astrocyte response to micromachined silicon implants,” Experimental Neurology 156, no.1 (1999): 33–49.

14. J. C. Williams, J. A. Hippensteel, J. Dilgen, W. Shain, and D. R. Kipke, “Complex impedance spectroscopy for monitoring tissue responses to inserted neural implants,” Journal of Neural Engineering 4, no. 4 (2007): 410–423. 10.1088/1741-2560/4/4/007

15. B. D. Ratner, “Reducing capsular thickness and enhancing angiogenesis around implant drug release systems,” Journal of Control Release 78, no. 1-3 (2002):211–8. doi: 10.1016/s0168-3659(01)00502-8. PMID: 11772462.

16. C. P. Millward, S. Perez da Rosa, D. Williams, G. Kokai, A. Byrne, B. Pettorini, “Foreign body granuloma secondary to ventriculo-peritoneal shunt: a rare scenario with a new insight,” Pediatric Neurosurgery 49, no. 4 (2013): 236–9. doi: 10.1159/000363330. Epub 2014 Jul 29. PMID: 25074235.

17. S. Gordon, “Alternative activation of macrophages,” Nature Reviews Immunology 3, no. 1 (2003): 23–35. 10.1038/nri978

18. D. A. Hume, I. L. Ross, S. R. Himes, R. T. Sasmono, C. A. Wells, and T. Ravasi, “The mononuclear phagocyte system revisited,” Journal of Leukocyte Biology 72, no.4(2002): 621–627. Retrieved from http://www.ncbi.nlm.nih.gov/pubmed/12377929

19. D. M. Mosser, “The many faces of macrophage activation,” Journal of Leukocyte Biology 73, no. 2 (2003): 209–212. 10.1189/jlb.0602325

20. D. M. Mosser, and J. P. Edwards, “Exploring the full spectrum of macrophage activation,” Nature Reviews Immunology 8, no. 12 (2008): 958–969. 10.1038/nri2448

21. B. Mytar, M. Siedlar, M. Woloszyn, I. Ruggiero, J. Pryjma, and M. Zembala, “Induction of reactive oxygen intermediates in human monocytes by tumour cells and their role in spontaneous monocyte cytotoxicity,” British Journal of Cancer 79, no. 5–6 (1999): 737–743. 10.1038/sj.bjc.6690118

22. T. M. Doherty, R. Kastelein, S. Menon, S. Andrade, and R. L. Coffman, “Modulation of murine macrophage function by IL-13,” Journal of Immunology 151, no. 12 (1993): 7151–7160. Retrieved from http://www.ncbi.nlm.nih.gov/pubmed/7903102

23. A. Gratchev, J. Kzhyshkowska, K. Köthe, et al., “Mphi1 and Mphi2 can be re-polarized by Th2 or Th1 cytokines, respectively, and respond to exogenous danger signals,” Immunobiology 211, no. 6–8 (2006): 473–486. 10.1016/j.imbio.2006.05.017

24. M. Munder, K. Eichmann, and M. Modolell, “Alternative metabolic states in murine macrophages reflected by the nitric oxide synthase/arginase balance: competitive regulation by CD4+ T cells correlates with Th1/Th2 phenotype,” Journal of Immunology 160, no. 11 (1998): 5347–5354. Retrieved from http://www.ncbi.nlm.nih.gov/pubmed/9605134

25. P. J. Murray, “Understanding and exploiting the endogenous interleukin-10/STAT3-mediated anti-inflammatory response,” Current Opinion in Pharmacology 6, no. 4 (2006): 379–386. 10.1016/j.coph.2006.01.010

26. M. Stein, S. Keshav, N. Harris, and S. Gordon, “Interleukin 4 potently enhances murine macrophage mannose receptor activity: a marker of alternative immunologic macrophage activation,” The Journal of Experimental Medicine 176, no. 1 (1992): 287–292. 10.1084/jem.176.1.287

27. F. A. W. Verreck, T. de Boer, D. M. L. Langenberg, et al., “Human IL-23-producing type 1 macrophages promote but IL-10-producing type 2 macrophages subvert immunity to (myco)bacteria,” Proceedings of the National Academy of Sciences of the United States of America 101, no. 13 (2004): 4560–4565. 10.1073/pnas.0400983101

28. M. Li, Q. Hou, L. Zhong, Y. Zhao, and X. Fu. “Macrophage related chronic inflammation in non-healing wounds,” Frontiers in Immunology 12 (2021): 681710. 10.3389/fimmu.2021.681710

29. M. Bartneck, K.-H. Heffels, Y. Pan, M. Bovi, G. Zwadlo-Klarwasser, and J. Groll, “Inducing healing-like human primary macrophage phenotypes by 3D hydrogel coated nanofibers,” Biomaterials 33, no. 16 (2012): 4136–4146. 10.1016/j.biomaterials.2012.02.050

30. R.-X. Wu, C. Ma, Y. Liang, F.-M. Chen, and X. Liu, “ECM-mimicking nanofibrous matrix coaxes macrophages toward an anti-inflammatory phenotype: Cellular behaviors and transcriptome analysis,” Applied materials today 18 (2020): 100508. doi: 10.1016/j.apmt.2019.100508

31. N. Li, S. Kang, Z. Liu, et al., “Immune-compatible designs of semiconducting polymers for bioelectronics with suppressed foreign-body response,” Nature materials 25, no. 1 (2026): 124–132. 10.1038/s41563-025-02213-x

32. X. Zhou, Z. Zhu, W. Dai, et al., “An immunocompatible conductive polymer for long-term bioelectronic implants,” Journal of the American Chemical Society 147, no. 42 (2025): 37985–37998. doi: 10.1021/jacs.5c07902. Epub 2025 Oct 9. PMID: 41069114.

33. S. F. Badylak, J. E. Valentin, A. K. Ravindra, G. P. McCabe, and A. M. Stewart-Akers, “Macrophage phenotype as a determinant of biologic scaffold remodeling,” Tissue Engineering Part A 14, no. 11 (2008): 1835–1842. 10.1089/ten.tea.2007.0264

34. B. N. Brown, B. D. Ratner, S. B. Goodman, S. Amar, and S. F. Badylak, “Macrophage polarization: an opportunity for improved outcomes in biomaterials and regenerative medicine,” Biomaterials 33, no. 15 (2012): 3792–3802. 10.1016/j.biomaterials.2012.02.034

35. B. N. Brown, J. E. Valentin, A. M. Stewart-Akers, G. P. McCabe, and S. F. Badylak, “Macrophage phenotype and remodeling outcomes in response to biologic scaffolds with and without a cellular component,” Biomaterials 30, no. 8 (2009): 1482–1491. 10.1016/j.biomaterials.2008.11.040

36. J. D., Bryers, C. M. Giachelli, and B. D. Ratner, “Engineering biomaterials to integrate and heal: the biocompatibility paradigm shifts,” Biotechnology and bioengineering 109, no. 8 (2012): 1898–1911. 10.1002/bit.24559

37. B. D. Ratner, and A. Marshall, “Porous biomaterials,” U.S. Patent 7,972,628, issued July 5, 2011. https://patents.google.com/patent/US7972628B2/en

38. S. Fili, P. Wölfelschneider, and M. Kohlhaas, “The STARflo glaucoma implant: preliminary 12 months results,” Graefe’s Archive for Clinical and Experimental Ophthalmology 256, no. 4 (2018): 773–781. 10.1007/s00417-018-3916-x

39. K. S., Thomson, F. S. Korte, C. M. Giachelli, B. D. Ratner, M. Regnier, and M. Scatena, “Prevascularized microtemplated fibrin scaffolds for cardiac tissue engineering applications,” Tissue Engineering Part A 19, no. 7-8 (2013): 967–977. 10.1089/ten.tea.2012.0286

40. E. M. Sussman, M. C. Halpin, J. Muster, R.T. Moon, and B. D. Ratner, “Porous implants modulate healing and induce shifts in local macrophage polarization in the foreign body reaction,” Annals of biomedical engineering 42, no. 7 (2014): 1508–1516. 10.1007/s10439-013-0933-0

41. T. F., Hady, B. Hwang, A. D. Pusic, R. L. Waworuntu, M. Mulligan, B. D. Ratner, and J. D. Bryers, “Uniform 40-µm-pore diameter precision templated scaffolds promote a pro-healing host response by extracellular vesicle immune communication,” Journal of tissue engineering and regenerative medicine 15, no. 1 (2021): 24–36. 10.1002/term.3160

42. L. Zhen, S. A. Creason, F. I. Simonovsky, et al., “Precision-porous polyurethane elastomers engineered for application in pro-healing vascular grafts: Synthesis, fabrication and detailed biocompatibility assessment,” Biomaterials 279 (2021): 121174.10.1016/j.biomaterials.2021.121174

43. N. R., Chan, B. Hwang, B. D. Ratner, and J. D. Bryers, “Monocytes contribute to a pro-healing response in 40 μm diameter uniform-pore, precision-templated scaffolds,” Journal of tissue engineering and regenerative medicine 16, no. 3 (2022): 297–310. 10.1002/term.3280

44. A. D., Bhrany, C. A. Irvin, K. Fujitani, Z. Liu, and B. D. Ratner. “Evaluation of a sphere-templated polymeric scaffold as a subcutaneous implant.” JAMA facial plastic surgery 15, no. 1 (2013): 29–33. 10.1001/2013.jamafacial.4

45. Y. Fukano, M. L. Usui, R. A. Underwood, et al., “Epidermal and dermal integration into sphere-templated porous poly (2-hydroxyethyl methacrylate) implants in mice,” Journal of Biomedical Materials Research Part A 94, no. 4 (2010): 1172–1186. 10.1002/jbm.a.32798

46. Y. Fukano, N. G. Knowles, M. L. Usui, et al., “Characterization of an in vitro model for evaluating the interface between skin and percutaneous biomaterials,” Wound repair and regeneration 14, no. 4 (2006): 484–491. 10.1111/j.1743-6109.2006.00138.x

47. S. N. Isenhath, Y. Fukano, M. L. Usui, et al., “A mouse model to evaluate the interface between skin and a percutaneous device,” Journal of Biomedical Materials Research Part A 83, no. 4 (2007): 915–922.10.1002/jbm.a.31391

48. N. G. Knowles, Y. Miyashita, M. L. Usui, et al., “A model for studying epithelial attachment and morphology at the interface between skin and percutaneous devices,” Journal of Biomedical Materials Research Part A 74, no. 3 (2005): 482–488. 10.1002/jbm.a.30384

49. R. A. Underwood, M. L. Usui, G. Zhao, et al., “Quantifying the effect of pore size and surface treatment on epidermal incorporation into percutaneously implanted sphere-templated porous biomaterials in mice,” Journal of Biomedical Materials Research Part A 98, no. 4 (2011): 499–508. 10.1002/jbm.a.33125

50. P. C. S. Bota, A. M. B. Collie, P. Puolakkainen, et al., “Biomaterial topography alters healing in vivo and monocyte/macrophage activation in vitro,” Journal of Biomedical Materials Research Part A 95, no. 2 (2010): 649–657. 10.1002/jbm.a.32893

51. A. Galperin, T. J. Long, S. Garty, and B. D. Ratner. “Synthesis and fabrication of a degradable poly (N-isopropyl acrylamide) scaffold for tissue engineering applications.” Journal of Biomedical Materials Research Part A 101, no. 3 (2013): 775–786. 10.1002/jbm.a.34380

52. S. K., Seidlits, Z. Z. Khaing, R. R. Petersen, et al., “The effects of hyaluronic acid hydrogels with tunable mechanical properties on neural progenitor cell differentiation,” Biomaterials 31, no. 14 (2010): 3930–3940. doi: 10.1016/j.biomaterials.2010.01.125. Epub 2010 Feb 19. PMID: 20171731.

53. J. P. Harris, A. E. Hess, S. J. Rowan, et al., “In vivo deployment of mechanically adaptive nanocomposites for intracortical microelectrodes,” Journal of neural engineering 8, no. 4 (2011): 046010. 10.1088/1741-2560/8/4/046010

54. J. K., Nguyen, D. J. Park, J. L. Skousen, et al., “Mechanically-compliant intracortical implants reduce the neuroinflammatory response,” Journal of neural engineering 11, no. 5 (2014): 056014. 10.1088/1741-2560/11/5/056014

55. W. Zhou, Y. Jiang, Q. Xu et al., “Soft and stretchable organic bioelectronics for continuous intraoperative neurophysiological monitoring during microsurgery,” Nature Biomedical Engineering 7, no. 10 (2023): 1270–1281. 10.1038/s41551-023-01069-3

56. L. Zhen, R. Darrow, N. Chen, et al., “Soft, precision engineered porous, hydrogel scaffolds mechanically tailored toward applications in the central nervous system,” Journal of Bioactive and Compatible Polymers 39, no. 6 (2024): 507–521. doi:10.1177/08839115241287215

57. J. Visser, F. P. W. Melchels, J. E. Jeon, et al., “Reinforcement of hydrogels using three-dimensionally printed microfibres,” Nature communications 6, no. 1 (2015): 6933.

58. S. Deitch, M. Choi, and P. Rousche. “Brain Model for Microelectrode Implantation Testing.” (2006).

59. F. Charlier, M. Weber, D. Izak, et al., “trevismd/statannotations: v0.5,*”* Zenodo (2022). 10.5281/zenodo.7213391

60. A. Ernst, K. Alkass, S. Bernard, et al., “Neurogenesis in the striatum of the adult human brain,” Cell 156, no. 5 (2014): 1072–1083.

61. M. V. Kokoeva, H. Yin, and J. S. Flier. “Evidence for constitutive neural cell proliferation in the adult murine hypothalamus.” Journal of Comparative Neurology 505, no. 2 (2007): 209–220.

62. K. C., Spencer, J. C. Sy, K. B. Ramadi, A. M. Graybiel, R. Langer, and M. J. Cima. “Characterization of mechanically matched hydrogel coatings to improve the biocompatibility of neural implants.” Scientific reports 7, no. 1 (2017): 1952.

63. L. R., Madden, D. J. Mortisen, E. M. Sussman, et al., “Proangiogenic scaffolds as functional templates for cardiac tissue engineering,” Proceedings of the National Academy of Sciences 107, no. 34 (2010): 15211–15216.

64. L. Zhen, E. Quiroga, S. A. Creason, et al., “Synthetic vascular graft that heals and regenerates,” Biomaterials 320 (2025): 123206.

65. K. L., Spiller, R. R. Anfang, K. J. Spiller, et al., “The role of macrophage phenotype in vascularization of tissue engineering scaffolds,” Biomaterials 35, no. 15 (2014): 4477–4488.

66. T. Stieglitz, H. Beutel, M. Schuettler, and J.-U. Meyer, “Micromachined, polyimide-based devices for flexible neural interfaces,” Biomedical microdevices 2, no. 4 (2000): 283–294.

67. K. Joseph, M. Kirsch, M. Johnston, et al., “Transcriptional characterization of the glial response due to chronic neural implantation of flexible microprobes,” Biomaterials 279 (2021): 121230.

68. S. Couillard-Despres, B. Winner, S. Schaubeck, et al., ”Doublecortin expression levels in adult brain reflect neurogenesis” European Journal of Neuroscience 21, no. 1 (2005): 1–14. 10.1111/j.1460-9568.2004.03813.x

69. Y. Chai, H. Zhao, S. Yang, et al. “Structural alignment guides oriented migration and differentiation of endogenous neural stem cells for neurogenesis in brain injury treatment.” Biomaterials 280 (2022): 121310.

70. K.-F. Huang, W.-C. Hsu, W.-T. Chiu, and J.-Y. Wang, “Functional improvement and neurogenesis after collagen-GAG matrix implantation into surgical brain trauma,” Biomaterials 33, no. 7 (2012): 2067–2075.

71. Y. Yang, Y. Fan, H. Zhang, et al., “Small molecules combined with collagen hydrogel direct neurogenesis and migration of neural stem cells after spinal cord injury,” Biomaterials 269 (2021): 120479.

72. J. Altman, and G. D. Das. “Autoradiographic and histological evidence of postnatal hippocampal neurogenesis in rats.” Journal of Comparative Neurology 124, no. 3 (1965): 319–335.

73. C. Lois, and A. Alvarez-Buylla, “Long-distance neuronal migration in the adult mammalian brain,” Science 264, no. 5162 (1994): 1145–1148.

74. E. Gould, A. J. Reeves, M. S. A. Graziano, and C. G. Gross, “Neurogenesis in the neocortex of adult primates,” Science 286, no. 5439 (1999): 548–552.

75. S. S., Magavi, B. R. Leavitt, and J. D. Macklis, “Induction of neurogenesis in the neocortex of adult mice,” Nature 405, no. 6789 (2000): 951–955.

76. W. Zheng, Q. ZhuGe, M. Zhong, et al., “Neurogenesis in adult human brain after traumatic brain injury.” Journal of neurotrauma 30, no. 22 (2013): 1872–1880.

77. B. Saha, S. Peron, K. Murray, M. Jaber, and A. Gaillard, “Cortical lesion stimulates adult subventricular zone neural progenitor cell proliferation and migration to the site of injury,” Stem cell research 11, no. 3 (2013): 965–977. 10.1016/j.scr.2013.06.006

78. J.-L., Yang, H. Fan, F.-F. Fu, et al., “Transient neurogenesis in ischemic cortex from Sox2+ astrocytes,” Neural Regeneration Research 18, no. 7 (2022): 1521.doi: 10.4103/1673-5374.357910. PMID: 36571357; PMCID: PMC10075105.

79. M. R. L. Dawson, A. Polito, J. M. Levine, and R. Reynolds. “NG2-expressing glial progenitor cells: an abundant and widespread population of cycling cells in the adult rat CNS.” Molecular and Cellular Neuroscience 24, no. 2 (2003): 476–488. doi: 10.1016/S1044-7431(03)00210-0

80. S. F., Bazarek, M. Thaqi, P. King, et al., “Engineered neurogenesis in naïve adult rat cortex by Ngn2-mediated neuronal reprogramming of resident oligodendrocyte progenitor cells,” Frontiers in Neuroscience 17 (2023): 1237176. 10.3389/fnins.2023.1237176

81. E. D., Laywell, P. Rakic, V. G. Kukekov, E. C. Holland, and D. A. Steindler, “Identification of a multipotent astrocytic stem cell in the immature and adult mouse brain,” Proceedings of the National Academy of Sciences 97, no. 25 (2000): 13883–13888.

82. T. Imura, H. I. Kornblum, and M. V. Sofroniew, “The predominant neural stem cell isolated from postnatal and adult forebrain but not early embryonic forebrain expresses GFAP,” Journal of Neuroscience 23, no. 7 (2003): 2824–2832.

83. Z. Guo, X. Wang, J. Xiao, et al., “Early postnatal GFAP-expressing cells produce multilineage progeny in cerebrum and astrocytes in cerebellum of adult mice,” Brain research 1532 (2013): 14–20.

84. W. Li, E. T. Mandeville, V. Durán-Laforet, et al., “Endothelial cells regulate astrocyte to neural progenitor cell trans-differentiation in a mouse model of stroke.” Nature communications 13, no. 1 (2022): 7812.

85. Y. Lei, X. Chen, J.-L. Mo, L.-L. Lv, Z.- W. Kou, and F.-Y. Sun, “Vascular endothelial growth factor promotes transdifferentiation of astrocytes into neurons via activation of the MAPK/Erk-Pax6 signal pathway,” Glia 71, no. 7 (2023): 1648–1666.

86. W.-R. Scha bitz, T. Steigleder, C. M. Cooper-Kuhn, et al., “Intravenous brain-derived neurotrophic factor enhances poststroke sensorimotor recovery and stimulates neurogenesis,” Stroke 38, no. 7 (2007): 2165–2172.

87. Y. Sun, K. Jin, L. Xie, et al., “VEGF-induced neuroprotection, neurogenesis, and angiogenesis after focal cerebral ischemia,” The Journal of clinical investigation 111, no. 12 (2003): 1843–1851.

88. D. H. Kim, Y. K. Seo, T. Thambi, et al., “Enhancing neurogenesis and angiogenesis with target delivery of stromal cell derived factor-1α using a dual ionic pH-sensitive copolymer.” Biomaterials 61 (2015): 115–125.

89. L. R. Nih, S. Gojgini, S. T. Carmichael, and T. Segura, “Dual-function injectable angiogenic biomaterial for the repair of brain tissue following stroke,” Nature materials 17, no. 7 (2018): 642–651.

90. P. Hao, H. Duan, F. Hao, et al., “Neural repair by NT3-chitosan via enhancement of endogenous neurogenesis after adult focal aspiration brain injury.” Biomaterials 140 (2017): 88–102.

91. Z. Álvarez, O. Castaño, A. A. Castells, et al., “Neurogenesis and vascularization of the damaged brain using a lactate-releasing biomimetic scaffold.” Biomaterials 35, no. 17 (2014): 4769–4781.

92. Y. Chang, B. Cho, E. Lee, et al., “Electromagnetized gold nanoparticles improve neurogenesis and cognition in the aged brain,” Biomaterials 278 (2021): 121157.

93. L. M. Crawford, Toward a Wholly Organic, Immunomodulatory Neuroelectronic Interface (Doctoral Dissertation), University of Washington, 2021.

