## Supplementary material for "Mechanically Compliant, Precision-Porous Brain Implants Reduce the Foreign Body Reaction and Guide Regeneration": Table S2

***Table S1*** *- Average stiffness values (kPa) for materials used in this study. Showing hydrated (left) and lyophilized (right) stiffness values.*

| **Water Content (%)** | **Nonporous** | | **100 µm pores** | | **40 µm pores** | |
| --- | --- | --- | --- | --- | --- | --- |
| **50** | 2,544 | 27,075 | 177 | 17,750 | 158 | 22,620 |
| **75** | 340 | 15,174 | 13.9 | 5,088 | 16.8 | 7,185 |
| **85** | 15.7 | 2,710 | 2 | 1,464 | 1.7 | 659 |

***Table S2*** *- Antibodies and immunohistochemistry information used in pHEMA/GMA implant experiments.*

| Antibody/stain | Target | Supplier | Catalog # | Host | Dilution |
| --- | --- | --- | --- | --- | --- |
| Hoechst 33342 | Cell Nuclei | ThermoFisher | H3570 | - | 1:1000 |
| Iba1 | Microglia | Wako | 016-20001 | Rabbit | 1:250 |
| GFAP | Astrocytes | ThermoFisher | 13-0300 | Rat | 1:800 |
| CD68 | Macrophages | Bio-Rad | MCA341R | Mouse | 1:500 |
| Arg1 | M2 Macrophages | Cell Signaling Technologies | 93668S | Rabbit | 1:100 |
| iNOS | M1 Macrophages | Abcam | ab15323 | Rabbit | 1:50 |
| NeuN | Neuronal Cell Bodies | Abcam | ab177487 | Rabbit | 1:500 |
| NeuroFilament | Neuronal Axons | Abcam | ab8135 | Rabbit | 1:1000 |
| MAP2 | Neurons/  dendrites | Millipore | ab5622 | Rabbit | 1:1000 |
| RECA1 | Endothelial Cells | Abcam | ab9774 | Mouse | 1:200 |
| Doublecortin | Neuronal Precursors | Santa Cruz | [E-6] sc-271390 | Mouse | 1:200 |
| AlexaFluor 488 Goat anti-Rabbit IgG (H+L) | Rabbit Primary Antibodies | ThermoFisher | A-11008 | Goat | 1:250 |
| AlexaFluor 594 Goat anti-Mouse IgG (H+L) | Mouse Primary Antibodies | ThermoFisher | A-11005 | Goat | 1:250 |
| AlexaFluor 594 Goat anti-Rat IgG (H+L) | Rat Primary Antibodies | ThermoFisher | A-11007 | Goat | 1:250 |

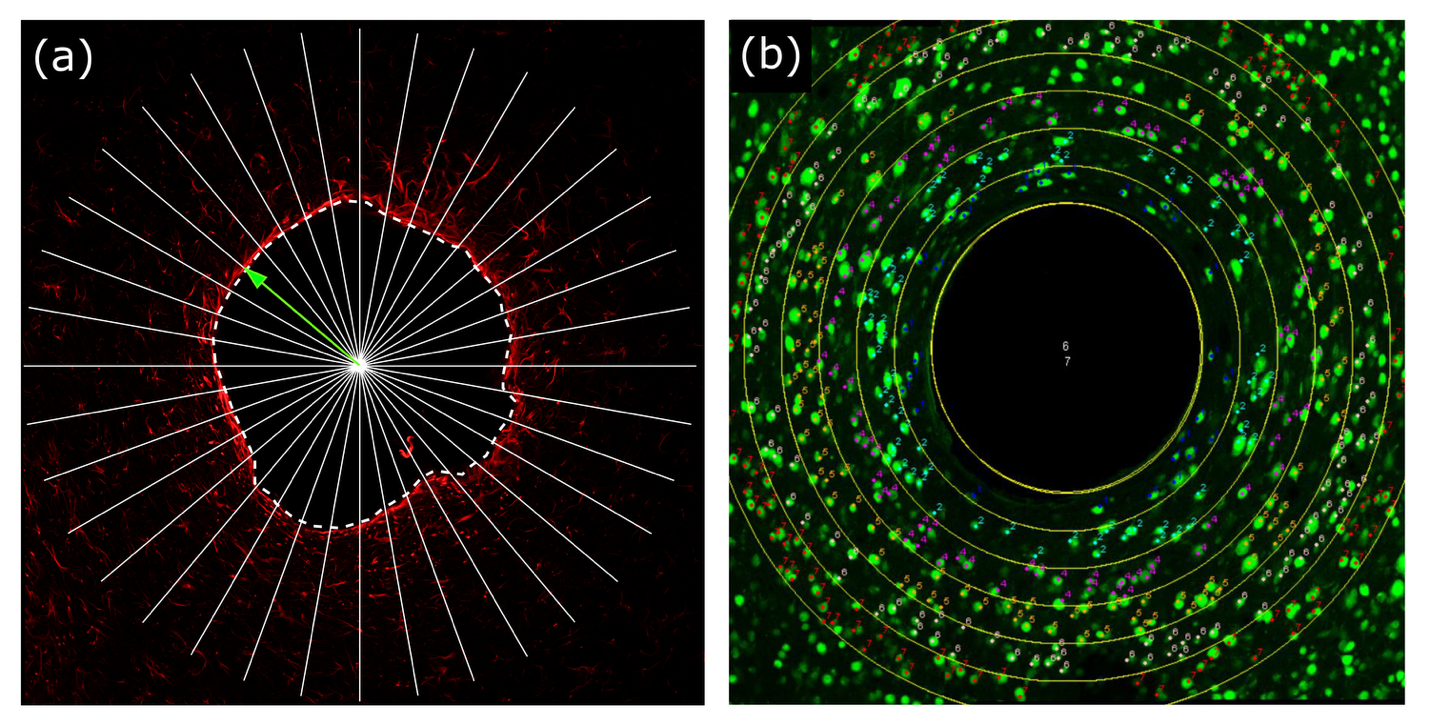

**Figure S1** – Examples of quantification measurements for two sample implants. **(a)** For fluorescence profile measurements used for GFAP and Iba1, the center point of each implant was selected, then fluorescence profile lines were captured by extending lines radially outwards every 10 degrees (for a total of 36 lines per implant), using the Radial Profile Angle plugin. For each line, the distance from the implant center to the implant edge (dashed line) was measured (green arrow), so fluorescence profiles could be shifted to start at the implant edge. Once aligned to the implant edge, all fluorescence profile lines from a given implant were averaged. The area under the averaged fluorescence profile from 0-25 μm away from the implant surface was calculated and used as the measure of fluorescence directly surrounding the implant. **(b)** For cell count measurements used for NeuN, CD68, Arg1, and iNOS, the outline of the implants were drawn. Then, 50 micron thick bands (NeuN) or 100 micron bands (CD68, Arg1, iNOS) were extended from the implant edge. The number of cells within each band was counted using the Cell Counter function in ImageJ and used for cell density calculations.
